# Ongoing activity in anatomically recurrent cortex is predominantly feedforward

**DOI:** 10.64898/2026.11.26.769545

**Authors:** Keshav Srinivasan, Tiago L. Ribeiro, Patrick Kells, Dietmar Plenz

## Abstract

The cerebral cortex is the major site of computation in the brain, and its recurrent wiring has motivated models in which repeated neuronal activation maintains information in stable attractor states. Large language models illustrate an alternative principle by transforming information layer by layer during each forward pass. A first step toward understanding what cortex and these models can teach us about each other is to establish whether feedforward dynamics can predominate in recurrent cortex. Here, we show that ongoing activity in the frontal cortex of awake mice is composed of parabolic neuronal avalanches in which activity progresses mainly through newly recruited neurons. Using two-photon calcium imaging in hundreds of simultaneously recorded layer II/III pyramidal neurons, we decomposed avalanche size into the number of participating neurons, mean activations per participant, and mean inferred spikes per activation. Avalanche size grew approximately quadratically with duration up to 0.5 s, primarily through recruitment of additional neurons. Participating neurons contributed close to one spike per avalanche on average. Newly recruited neurons also dominated the rise and fall of the average avalanche profile. An unbiased recruitment model, in which all neurons have the same selection probability, accounted for the predominance of new over repeat participation, showing that feedforward recruitment can dominate even when previously active neurons remain available for reactivation. Parabolic avalanches thus express predominantly feedforward dynamics within anatomically recurrent cortex. Cortical computation may therefore have more in common with feedforward processing in large language models than recurrent anatomy alone would suggest.

## Introduction

How computation emerges from the collective activity of neurons is a central question in neuroscience [1]. In the cerebral cortex, recurrent connections allow activity to return to neurons that have already fired [2,3]. In layers II/III, these circuits include neighboring pyramidal neurons that excite each other directly [4,5]. Such recurrence supports attractor models of associative memory [6], but recurrent anatomy does not determine how strongly neurons are reactivated during individual population events. A recurrent network could express predominantly feedforward dynamics if activity progresses mainly through newly recruited neurons, with limited return to previous participants. Abeles proposed that synchronous spikes could propagate from one group of cortical neurons to the next along feedforward connections, an arrangement he called a synfire chain [7,8]. Establishing whether this organization occurs in awake cortex would provide an experimental basis for connecting cortical dynamics to feedforward processing in artificial networks.

Feedforward artificial neural networks implement computation by transforming representations through successive layers [9]. Transformer-based large language models likewise process information through successive attention and feedforward layers within each forward pass [10,11]. The success of these systems raises a fundamental question about the brain: how does cortical activity organize the progression through neuronal populations that could support successive stages of processing?

Neuronal avalanches are scale-invariant cascades of cortical activity [12,13] observed in awake rodents [14,15] and nonhuman primates [16]. Their scaling links cortical activity to the physics of criticality [12]. Local field potential recordings [16] showed that avalanches in the scaling regime rarely revisited the same sites [17], and single-neuron recordings showed that individual neurons participated selectively in particular avalanche patterns [18]. By tracking individual neurons with calcium imaging [15], we previously showed that mean avalanche size in superficial cortex grows approximately with the square of its duration [19]. Does this growth arise mainly from recruiting new neurons, as expected for feedforward dynamics, or from reactivating earlier participants, as expected for recurrent dynamics?

As avalanches unfolded in awake mouse frontal cortex, activity progressed through successive groups of neurons and rarely returned to those already active. A model that gave every neuron the same chance of selection predicted the balance of first and repeat activations, showing how feedforward recruitment can dominate even when earlier participants remain available. We conclude that parabolic avalanches represent a predominantly feedforward organization of ongoing activity in anatomically recurrent cortex.

## Results

### Decomposition separates recruitment from repeat activation

Population activity can grow through recruitment of additional neurons, repeated activation of previous participants, or greater activity within each activation. To distinguish these contributions, we analyzed ongoing activity recorded by two-photon calcium imaging in the anterior cingulate cortex (ACC) and medial prefrontal cortex (mPFC) of awake C57BL/6 mice expressing the genetically encoded calcium indicator jRGECO1a after viral injection as reported previously [19]. A prism allowed for the contralateral recording of these medially positioned cortical areas across the midline. The archived dataset comprised 17 recordings from 5 mice, with 138–321 neurons per recording (Fig. 1a). Building on our demonstration of parabolic neuronal avalanches in these recordings [19], we tracked neuronal identity throughout each event.

**Figure 1.**
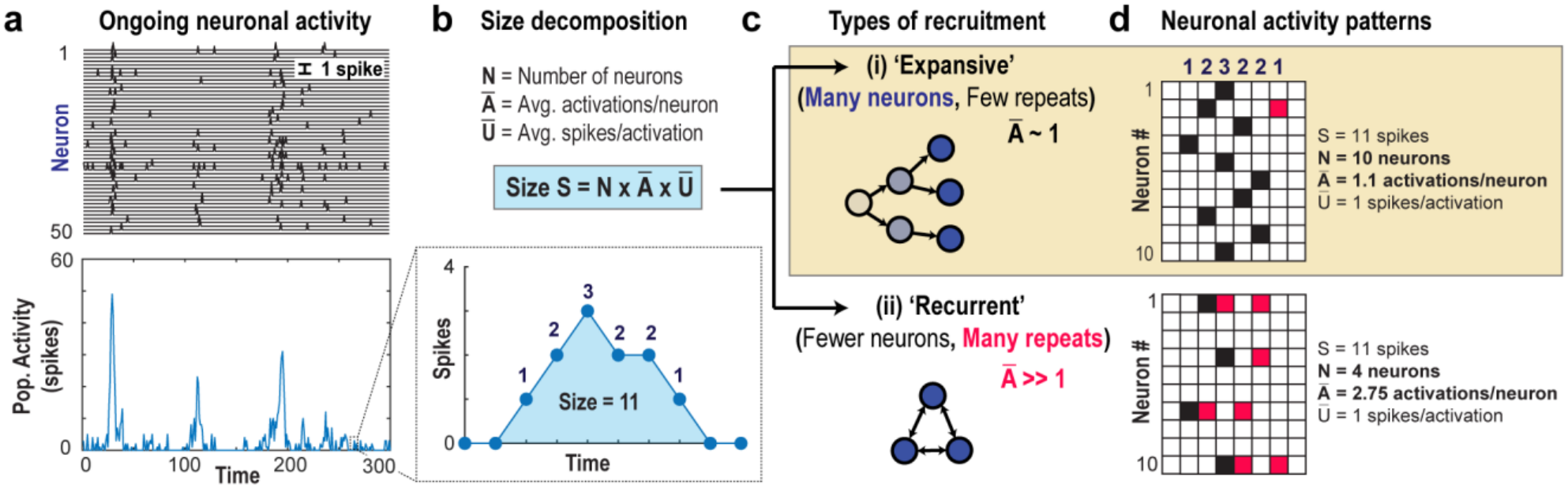
Avalanche decomposition distinguishes recruitment from repeated participation. a, Example thresholded activity from 50 neurons in the anterior cingulate cortex of awake mice expressing jRGECO1a (top), with the corresponding population activity (bottom) [19]. b, Decomposition of avalanche size into the number of participating neurons, N; mean activations per participating neuron, Ā; and mean activity per activation, Ū. The illustrated avalanche spans 6 active bins and has size 11. c, Schematic recruitment patterns: engagement of additional neurons with limited repeat participation, and repeated participation of a smaller neuronal group. The schematics illustrate possible patterns and do not represent measured synaptic connectivity. d, Two illustrative rasters with the same size, duration and population temporal profile but different neuronal participation. The upper raster has N = 10, Ā = 1.1 and Ū = 1; the lower raster has N = 4, Ā = 2.75 and Ū = 1. Repeat activations are highlighted in red. An activation is a positive contribution by one neuron in one analysis bin after population-based thresholding; activity is inferred from calcium fluorescence and subsequent spike deconvolution.

For each avalanche, we counted distinct participating neurons, N, and occupied neuron–time bins, B. An activation denotes a positive contribution by one neuron within one analysis bin after population-based thresholding. Mean activations per participant are *Ā* = *B*/*N*, and mean inferred activity per activation is *Ū* = *S*/*B*. Here, S denotes avalanche size in inferred spikes, giving the exact event-level identity *S* = *N* × *Ā* × *Ū* (Fig. 1b). Two avalanches can therefore have the same size and population temporal profile while recruiting different numbers of neurons (Fig. 1c,d). In the matched examples, limited repeat participation accompanies recruitment of a larger group, whereas frequent repeats engage a smaller group. This decomposition identifies the organization of detected participation; it does not reconstruct the synaptic paths between neurons.

### Short-duration avalanches recover near-quadratic scaling

Spatial subsampling can fragment a population event by missing consecutive active neurons, which can be overcome by temporal coarse-graining if the system is critical [20]. Accordingly, we first examined how the size– duration relationship changed with temporal coarse-graining. The factor *k* specifies the number of original imaging frames combined into an analysis bin. Across *k* = 1–25, the estimated exponent depended on both bin width and the fitted duration interval (Fig. 2a,b). At *k* = 6, corresponding to approximately 132 ms per bin, mean avalanche size grew with exponent χ = 1.96 (95% confidence interval, 1.71–2.22) over avalanche durations *L* = 1–4 bins, or approximately 0.13–0.53 s. This recovers the near-quadratic relationship reported previously [19], consistent with the recovery of parabolic scaling under temporal coarse-graining in spatially subsampled critical networks [20].

**Figure 2.**
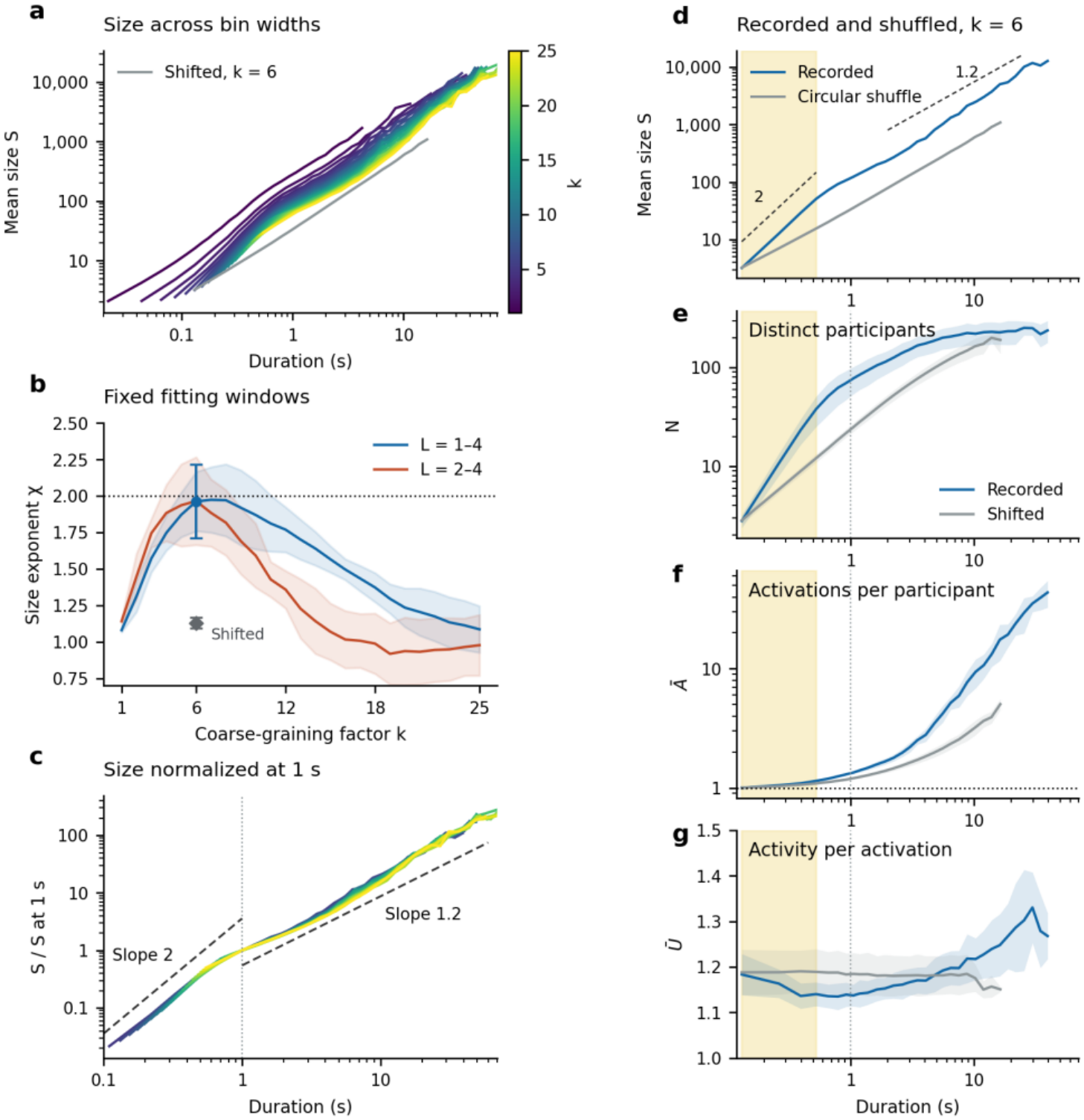
Recruitment of additional neurons dominates short-duration avalanche growth. a, Mean size S versus duration for k = 1–25 (color key); gray, circular-shift control at k = 6. b, Size–duration exponents fitted to exact duration-conditioned means over L = 1–4 (blue) and L = 2–4 (red). L denotes duration in analysis bins; the physical interval therefore changes with k. At k = 6, the blue circle and gray diamond show four-duration estimates for recorded and circularly shifted activity, respectively. Error bars show 95% confidence intervals across five mice. Fits use unweighted ordinary least squares in log–log coordinates. The dotted horizontal line is χ = 2, a reference rather than a fitted value. c, Mean size for k = 5–25 divided by its value at 1 s, obtained by interpolation in log–log coordinates. This is a vertical normalization; the duration axis is not rescaled. Dashed lines mark reference slopes of 2 (0.1–1 s) and 1.2 (1–60 s). d, Mean avalanche size versus duration at k = 6 for recorded activity (blue) and the circular-shift control (gray). Dashed lines indicate reference slopes of 2 and 1.2. e–g, N, *Ā* and *Ū* at k = 6 for recorded activity (blue) and independently shifted neuronal traces (gray; five shifts per recording with thresholds refitted). Yellow shading in d–g marks the short scaling-fit interval, L = 1–4 at k = 6 (approximately 0.13–0.53 s); the dotted vertical line in e–g marks the 1 s upper limit used for multibin participation summaries. Numerical multibin summaries exclude L = 1. In f, the dotted line is *Ā* = 1; one-bin avalanches lie on this line by definition. Duration curves show exact durations through L = 9 and fixed logarithmic bins thereafter, plotted at the mean observed duration within each bin. At least 10 event views are required per recording and displayed duration group, with coverage by at least three mice. Fits use exact durations, not the logarithmically binned display. Data in Figures 2–4 comprise 17 recordings from 5 mice. Recording-level estimates were averaged within each mouse, then mice were weighted equally. Bands show one standard deviation across mice; numerical confidence intervals in the Results are 95% t intervals across the five mouse estimates. Temporal offsets provide overlapping views of events, not independent biological samples.

Excluding one-bin avalanches gave χ = 1.97 (95% confidence interval, 1.59–2.34) over L = 2–4. Curves for k = 5–25 also approached a common relationship after their mean sizes were normalized at 1 s (Fig. 2c), extending the comparison with previous recordings [19] and subsampling analyses [20]. This normalization does not establish quadratic scaling throughout the displayed range. The departure from near-quadratic growth beyond approximately 0.5 s is consistent with the finite spatial observation window. We distinguish this short scaling-fit interval from the broader interval used to quantify neuronal participation. Beyond 10 s, avalanches were too sparsely sampled to estimate the long-duration scaling exponent reliably. At k = 6, the number of contributing recordings fell from 15 at 10.3 s to three at 39.4 s.

To test whether near-quadratic growth required temporal coordination between neurons, we independently circularly shifted their traces and repeated the thresholding and avalanche analysis (Fig. 2d). This preserves the time course within each trace while disrupting its temporal relation to other neurons. Across five shifts per recording, the exponent at *k* = 6 and *L* = 1–4 was 1.13 (95% confidence interval, 1.09–1.17) (Fig. 2b). The paired difference between recorded and shifted activity was 0.83 (95% confidence interval, 0.57–1.09) across mice. The coordinated short-duration regime is therefore distinct from the shallower relationship in the shifted control. Long-duration events must be examined separately rather than used to characterize this short-duration regime.

### Near-quadratic growth reflects feedforward recruitment

We next asked which component of avalanche size accounts for its steep increase with duration. Over the same L = 1–4 interval used for the size exponent at k = 6, distinct neuronal participation grew with exponent 1.86 (95% confidence interval, 1.60–2.11) (Fig. 2e). The corresponding exponents for Ā and Ū were 0.093 and −0.030, respectively (Fig. 2f,g). Thus, additional neurons joining the avalanche accounted for nearly all of the near-quadratic growth.

A mean activation count near one does not specify how many neurons participate only once, and one-bin avalanches have Ā = 1 by definition. We therefore quantified repeat participation separately in multibin avalanches lasting up to 1 s. Mean Ā was 1.147 (95% confidence interval, 1.130–1.163). The fraction of participants active in exactly one bin averaged 87.6% (95% confidence interval, 86.3–89.0%) across avalanches, and repeat activations accounted for 11.5% (95% confidence interval, 10.5–12.5%) of occupied neuron–time bins. These measurements establish predominantly feedforward recruitment at the analysis-bin scale: most participating neurons contribute in one bin, while repeat participation is present but modest. Multiple inferred spikes within a bin contribute to Ū and are not counted as separate activations. Mean Ū was 1.144 (95% confidence interval, 1.110–1.179).

For longer avalanches, recruitment became shallower as repeat participation increased (Fig. 2e,f). Over the broader *L* = 2–7 interval, the recruitment exponent was 1.50 (95% confidence interval, 1.28–1.71), below its estimate in the shorter scaling-fit interval. At still longer durations, the number of recruited neurons approached a plateau set by the recorded population, whereas Ā continued to increase. The transition is therefore gradual. The observed field of view constrains the measured recruitment pool; neurons outside it remain unobserved.

We asked whether the neurons most strongly correlated with the rest of the population better preserved avalanche scaling than an equally sized random subset (Fig. 3a). Correlation-ranked subsets showed steeper size– duration relationships at some bin widths when the same data were used for ranking and evaluation. When we ranked neurons on one half of each recording and tested on the other, the exponent fitted across four durations (0.26–1.05 s) at 20% retention and k = 12 averaged 1.58 for correlation-ranked subsets and 1.48 for random subsets (paired difference 0.10, 95% confidence interval −0.12 to 0.32, two-sided paired t test across five mice, t(4) = 1.25, P = 0.28). In this held-out comparison, we found no clear advantage to selecting neurons by their correlation with population activity.

**Figure 3.**
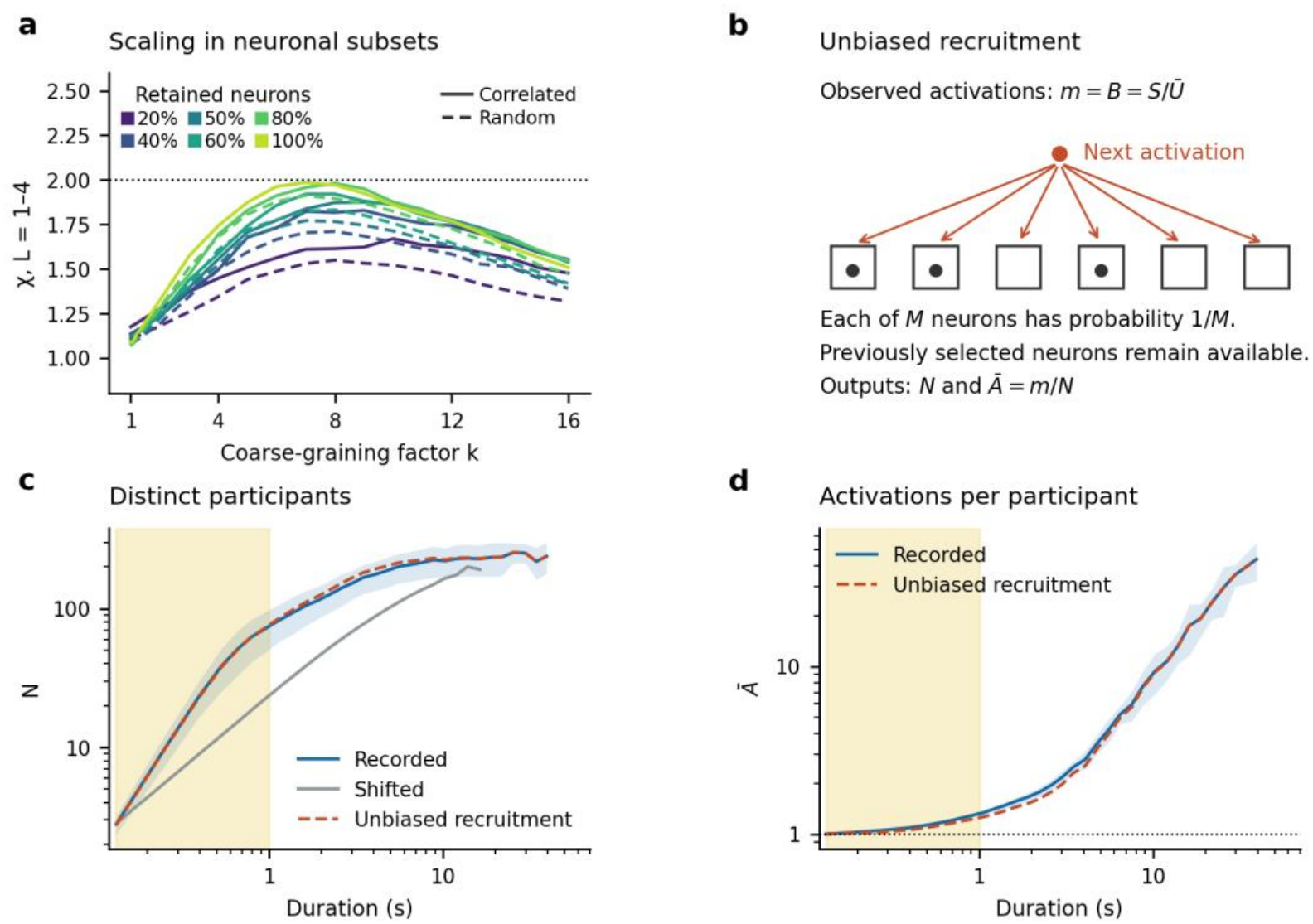
Subset selection affects scaling, and unbiased recruitment predicts the main participation pattern. a, Size–duration exponent χ fitted over L = 1–4 bins versus k. Color indicates the fraction of recorded neurons retained. Solid lines show subsets ranked by Pearson correlation with the activity of the other neurons in the same recording. Dashed lines show the mean of five nested random selections per recording. At 100% retention the two curves coincide. Thresholds are re-estimated for each retained subset and k. The dotted line is χ = 2. Curves show mouse means; held-out selection and its uncertainty are quantified in the Results. b, Schematic of unbiased recruitment. Each of m observed activations selects one of M recorded neurons with probability 1/M. Black dots mark previous selections; these neurons remain available for the next activation. The model predicts distinct participants, N, and mean activations per participant, Ā = m/N. c, Distinct participants at k = 6 in recorded activity (blue), shifted activity (gray), and the unbiased model (red dashed). d, Recorded Ā (blue) and its exact model expectation 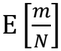 (red dashed). For each event, m is the observed number of occupied neuron–time bins. Model predictions are computed per event before averaging and have no fitted pool parameter. Yellow shading in c,d marks durations from one analysis bin to 1 s; numerical participation summaries exclude one-bin events. Duration-bin conventions follow Fig. 2. Only recorded-data uncertainty is shaded in c,d. Alternative recruitment models and the ratio approximation for Ā are shown in Supplementary Fig. 1. Data in Figures 2–4 comprise 17 recordings from 5 mice. Recording-level estimates were averaged within each mouse, then mice were weighted equally. Bands show one standard deviation across mice; numerical confidence intervals in the Results are 95% t intervals across the five mouse estimates. Temporal offsets provide overlapping views of events, not independent biological samples.

### An unbiased recruitment model accounts for most of the participation pattern

Does recruitment dominance require a rule that preferentially selects neurons not yet engaged in an avalanche? We tested an unbiased recruitment model in which every neuron has the same selection probability and previous participants remain available for reactivation. For each recorded avalanche, the model assigns 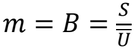 activations independently to a pool of M neurons, with M fixed to the number of neurons recorded in that session (Fig. 3b). Its expected number of distinct participants is 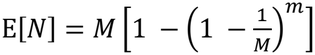. No pool size was fitted. The model therefore predicts participation conditional on observed activity, rather than generating the activity itself (Fig. 3c,d).

Unbiased recruitment followed the overall increase in N and the rise in Ā. For multibin avalanches lasting up to 1 s at k = 6, it overpredicted N by an average of 4.1% (95% confidence interval, 2.1–6.1%), with a mean absolute relative error of 7.5% (95% confidence interval, 6.6–8.4%). Predictions were computed for individual avalanches before averaging. Two comparison models tested whether preserving the active-neuron count within each bin or accounting for unequal neuronal activity rates improved this agreement. Neither reduced the mean absolute relative error in N (Supplementary Fig. 1a). For Ā, we used the exact model expectation 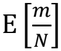 the closely overlapping ratio 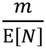 bis shown separately in Supplementary Fig. 1b.

### First activations dominate the avalanche profile

Total recruitment does not reveal when neurons first enter an avalanche. We therefore separated the temporal profiles of first and repeat participation for *L* = 2–7 at *k* = 6 (Fig. 4a–e). In profiles with three or more bins, mean population activity rose and fell with the number of simultaneously active neurons, whereas inferred activity per active neuron varied less. These profiles use only observed active bins; empty start and endpoints were not inserted to impose a profile shape.

**Figure 4.**
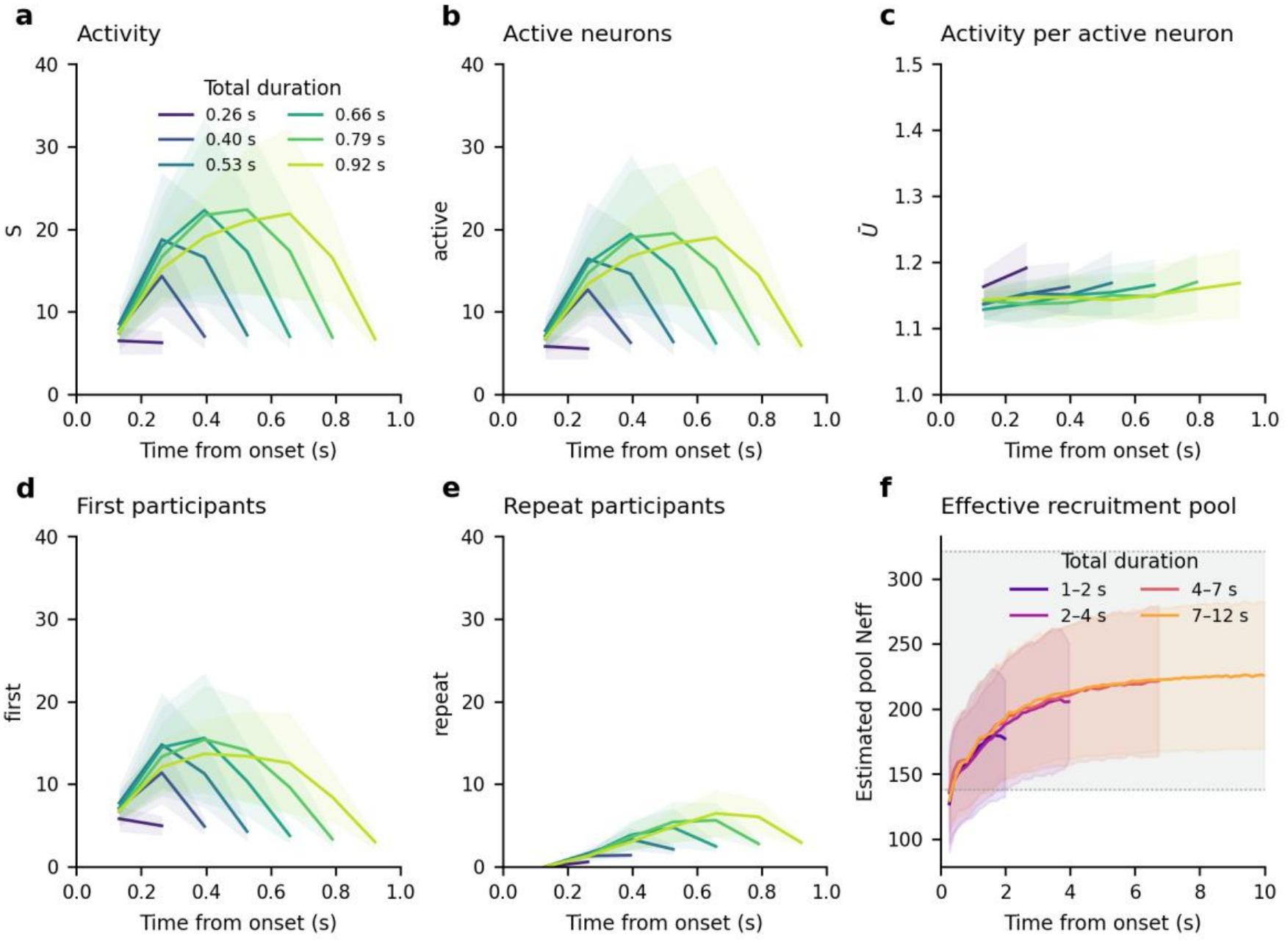
First participation dominates the temporal profile of short avalanches. a–e, Temporal profiles for exact durations L = 2–7 at k = 6, corresponding to approximately 0.26, 0.40, 0.53, 0.66, 0.79 and 0.92 s (color key in a). a, Inferred population activity per bin. b, Number of neurons active in that bin, distinct from N for the whole avalanche. c, Inferred activity per active neuron. d, Neurons active for the first time in that avalanche. e, Neurons active after participation in an earlier bin. Panels a, b, d and e share a y-axis range of 0–40 for absolute comparison; panel a reports inferred activity, whereas b, d and e report neuron counts. First and repeat counts sum exactly to the active-neuron count for each event and bin. Profiles are aligned to onset and displayed at bin ends without artificial zero endpoints. At least 10 event views are required per recording and exact duration. f, Effective recruitment pool for total durations 1–2, 2–4, 4–7 and 7–12 s (lower bounds included, upper bounds excluded). Within each recording, elapsed bin and duration group, 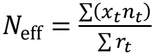, where x_t_ counts distinct neurons active before bin t, n_t_ counts currently active neurons and r_t_ counts current repeat participants. Sums run over event views. At least 10 views are required; zero-repeat estimates are undefined and omitted. Estimates are not capped at the recorded neuronal count. The gray band marks the range of recorded neurons across sessions (138–321), not uncertainty in the pool estimate. The displayed range ends at 10 s; points require at least three contributing mice. This estimator measures accessibility under uniform selection, not anatomical pool size. Data in Figures 2–4 comprise 17 recordings from 5 mice. Recording-level estimates were averaged within each mouse, then mice were weighted equally. Bands show one standard deviation across mice; numerical confidence intervals in the Results are 95% t intervals across the five mouse estimates. Temporal offsets provide overlapping views of events, not independent biological samples.

First participation exceeded repeat participation at every displayed bin in the mouse-averaged profiles (Fig. 4d,e). Repeats accumulated later, approaching the first-participation count near the end of the longest illustrated events. The decomposition thus connects population growth and its temporal profile to the same cellular organization: newly recruited neurons dominate the event, while returns to previous participants contribute increasingly as the event proceeds. This supports predominantly feedforward dynamics at the analysis-bin scale without requiring strictly single activation of every neuron.

### Repeat probability estimates an effective recruitment pool

Repeat participation can also be used to infer an effective accessible pool from the data. In the model above, the recorded pool size predicts repeat probability. Here, we inverted that relationship: under unbiased sampling, x previous participants occupy a fraction 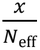 of an accessible pool of *N*_eff_ neurons. We estimated *N*_eff_ from the summed products of prior distinct participants and current active-neuron counts, divided by the summed current repeat counts, within each recording, elapsed bin and duration group (Fig. 4f). Previous participants were counted only from earlier bins. Estimates with no observed repeats were left undefined.

The estimated pool increased with elapsed time, with similar trajectories across the displayed duration groups of 1–2, 2–4, 4–7 and 7–12 s. For context, the recordings contained 138–321 neurons; this range is shown in Fig. 4f.

The estimates describe effective accessibility under the equal-selection assumption, rather than the number of neurons directly connected to each participant. Rare repeats make early estimates sensitive to the averaging rule.

## Discussion

### Feedforward organization within recurrent cortex

Recurrent cortical connections permit feedback, but does ongoing activity repeatedly engage the same neurons, or does it progress mainly through newly recruited neurons in a feedforward sequence? We demonstrated that parabolic neuronal avalanches in awake mouse frontal cortex grow predominantly by recruiting additional neurons. In multibin avalanches lasting up to 1 s, an average of 87.6% of participants were active in exactly one analysis bin, and first participation dominated the temporal profile. These findings identify a predominantly feedforward organization of neuronal participation within recurrent cortical anatomy.

This organization gives the comparison between biological and artificial computation an experimental basis. Feedforward artificial networks transform representations through successive layers [9], including the attention and feedforward layers traversed during a forward pass in transformer-based large language models [10,11]. Cortical avalanches exhibit a progression through successive neuronal groups within recurrent circuitry. A neuron’s position in this progression is defined by when it participates in the event. The correspondence concerns the organization of activity during an event or forward pass; autoregressive generation also reuses previous outputs across passes. Whether successive recruited groups transform information as an avalanche unfolds is a concrete experimental question arising from this comparison.

Synfire chains provide a specific model for realizing feedforward propagation through predefined synaptic pathways [7,8]. Convergent and divergent connections link successive neuronal groups and permit the stable propagation of synchronous spike volleys [21]. Embedding such pathways in recurrent networks can recruit excessive background excitation [22]; appropriately sized inhibitory pools accompanying the excitatory groups can stabilize propagation through “shadow inhibition” [23]. These models specify the progression through neuronal groups in their connectivity.

Hemberger et al. provided direct experimental evidence for chained synaptic activation in turtle cortex: brief stimulation of individual pyramidal neurons triggered shortlasting (<200 ms), reliable sequences involving excitatory and inhibitory neurons [24]. Their findings establish preferred functional pathways through which activity can propagate without uncontrolled amplification. Our measurements address spontaneous, parabolically scaling activity in awake mammalian cortex, identifying predominantly feedforward participation without specifying a chain in advance.

Experimentally, it is well established that avalanches are embedded in a scale-invariant hierarchy of avalanches within avalanches. We first demonstrated this nesting in awake monkey cortex: the power-law distribution of avalanche sizes remained unchanged when the amplitude threshold for detecting negative local field potential events was varied [16]. Lowering the threshold revealed smaller-amplitude avalanches within the apparently quiet intervals between larger-amplitude avalanches, preserving the same statistical organization across scales [16]. Across several millimeters of awake macaque cortex, Yu et al. showed that negative local field potential peaks rarely recurred at the same site within an avalanche in the power-law regime, establishing the predominantly feedforward character of avalanche dynamics at the mesoscale [17]. Within this nested organization, the same feedforward pattern emerges at both scales as avalanches recruit mainly new neurons in our recordings and mainly new sites of local population activity in the macaque recordings. In cortical slice cultures, we subsequently quantified the relationships between avalanche size, intervening quiet times, and subsequent avalanches, and established their dependence on excitation–inhibition balance [25,26,27]. The resulting picture is one in which “avalanches beget avalanches” [12].

This temporal organization suggests that feedforward propagation can be stacked across events. Within an avalanche, activity advances predominantly through successive neuronal groups; across avalanches, the activity and network state left by one event can shape the next. Successive avalanches could thus concatenate episodes of layered propagation into longer sequences within recurrent cortex. This extends the computational perspective from the internal organization of a single cascade to the composition of multiple cascades. Establishing how neuronal participation and information are transferred between successive avalanches would test whether this stacking supports multistage computation.

### Cellular organization of avalanche scaling

The decomposition *S* = *N × Ā × Ū* connects a collective scaling law to its cellular constituents: the number of participating neurons, their mean number of activations, and the mean inferred activity per activation. Quadratic growth could, in principle, be carried by any combination of these contributions. We found that recruitment provides the dominant contribution, while first participation also shapes the evolving temporal profile. The scaling of population activity therefore has a specific cellular expression: an avalanche grows mainly by engaging additional neurons, with limited repeated activation of those already engaged.

This conclusion depends on the regime in which scaling is resolved. At approximately 132 ms per analysis bin, the size–duration exponent was 1.96 over approximately 0.13–0.53 s, compared with 1.13 after independently shifting neuronal traces in time. The control preserves individual activity traces while disrupting their temporal coordination, linking near-quadratic growth to coordinated population activity. The above upper duration limit reflects a finite-size effect imposed by the spatial observation window. Using a mesoscope to image individual neurons over approximately 1 mm^2^ of mouse somatosensory cortex, we previously found that parabolic scaling extended to avalanches lasting about 5 s [19]. In the smaller frontal cortical field studied here, the number of participating neurons grew less steeply with avalanche duration, and repeat participation increased in longer events. The interval supporting near-quadratic scaling must therefore be distinguished from the broader interval used to measure participation.

In the macaque recordings, an avalanche with more negative local field potential peaks than recording sites must revisit at least one site, regardless of which neurons contribute to each peak [17]. Using avalanches reconstructed from inferred functional connections, Salners et al. found that most small avalanches were feedforward, with occasional cycles, whereas large avalanches beyond the power-law regime exhibited pronounced recurrent and cyclic activity [28]. These observations reinforce the need to distinguish dynamical regimes. Recurrence in the distribution tail cannot by itself establish that repeated neuronal activation drives parabolic avalanche growth.

### A statistical basis for recruitment dominance

The unbiased recruitment model shows how predominantly new participation can arise even when every previous participant remains available for selection. When relatively few activations are distributed across a larger pool, most reach neurons that have not yet participated. Equal selection therefore accounts for much of the observed participation pattern without an explicit rule that avoids previous participants. This provides a simple statistical interpretation of recruitment dominance, without requiring that cortical neurons have identical activity rates.

The model isolates neuronal occupancy from the dynamics that generate temporal coordination and parabolic scaling. Because it receives the observed activation count as input, it tests the balance of recruitment and repetition without specifying the sequence of propagation. We fixed its pool size to the recorded neuronal count and compared equal selection with rate-weighted and within-bin sampling. For longer events, the observed repeat probability yielded an effective pool that increased with elapsed time. This is consistent with an avalanche progressively reaching a larger set of neurons through successive interactions, without requiring direct connections between all participants. The estimate remains conditional on equal selection and the estimation of repeat probability. The finite field of view and unobserved neurons prevent interpreting it as a direct anatomical count.

### Flexible assemblies and cortical computation

We found that avalanches grow mainly by recruiting new neurons, raising the question of whether it matters which neurons join. Manley et al. showed that recording more cortical neurons reveals activity patterns that smaller samples miss [29]. Rigotti et al. showed why such patterns can matter for behavior: prefrontal neurons responded to combinations of task features, and the dimensionality of population activity fell on error trials [30]. Recruitment thus offers a possible link to cortical representation through the different combinations of neurons that can participate in avalanches of a given size.

Brief participation within one avalanche also permits a neuron to contribute to many successive events. Its average firing rate could change through how often it participates, even when its contribution to each event remains brief. This provides a temporal setting for flexible assembly membership. Prefrontal neurons contribute differently to perceptual and mnemonic representations [31]; avalanche-resolved analysis could test whether these differences are expressed through changes in the groups a neuron joins across events.

Temporal multiplexing offers a related test. Single neurons can alternate between response patterns associated with simultaneously presented stimuli [32], with coordinated fluctuations observed in visual cortex and other visual processing regions [33,34]. Determining whether these representations occupy successive avalanches, or successive recruited groups within an avalanche, would connect the participation pattern to information processing. This distinction also clarifies why brief coordinated events need not imply large trial-to-trial spike-count correlations [35]: the temporal window and definition of correlation determine which organization is measured.

### Measurement scope and outlook

The cellular interpretation is set by the sensitivity and temporal resolution of the recordings. In our recordings, an activation denotes detected neuronal activity in one analysis bin, with minimal activity events corresponding to approximately 1–2 spikes given the sensitivity of jRGECO1a [36]. Calcium imaging and temporal coarse-graining can combine several action potentials within one inferred activation or occupied bin. Such activity contributes to *Ū. Ā* includes the first activation and increases with repeat participation across bins. A value of *Ā* near one therefore describes participation across bins, and a value of *Ū* near one requires calibration before it can be interpreted as one action potential. Thresholding and spatial subsampling can further alter the observed balance of first and repeat participation. Electrical recordings and targeted perturbations could resolve within-bin firing and test causal transfer between successive groups, extending the present measurements of participation to the interactions that generate it.

Parabolic avalanches reveal that recurrent cortical networks can organize activity predominantly through successive neuronal recruitment. This establishes a measurable connection between collective cortical dynamics and feedforward processing in large language models, and a basis for testing how neuronal cascades support successive stages of processing.

## Methods

### Animals and two-photon calcium imaging

All experiments were approved by the Animal Care and Use Committee of the National Institute of Mental Health. C57BL/6 mice were injected with a viral construct expressing jRGECO1a under a synapsin promoter. Chronic two-photon imaging was performed in the anterior cingulate and medial prefrontal cortex through a microprism, as described previously [19]. The present analysis used all 17 recording entries in the five archived jRGECO-deepIP animal files, comprising 138–321 neurons per recording. All recorded neurons were included in the full-population analysis. Recordings were analyzed separately before aggregation within mice.

Images were acquired with a scanning microscope (Bergamo II series, B248, Thorlabs) coupled to a pulsed femtosecond two-photon laser (Chameleon Discovery NX, Coherent). The excitation wavelength was 1120 nm. The field of view was approximately 450 μm × 450 μm. Images of 512 × 512 pixels were acquired at 45.527 Hz by bidirectional scanning with a 12 kHz galvo-resonant scanner. Average power at the sample was <70 mW.

Imaging movies stored as unsigned 16-bit TIFF files underwent rigid motion correction in suite2p [37], denoising with DeepInterpolation [38], and semiautomatic region-of-interest selection and fluorescence extraction in suite2p. For each neuronal region of interest, the corrected fluorescence was *F*_corrected_ = *F*_ROI_ − 0.7 × *F*_neuropil_, where F_ROI_ is somatic fluorescence and F_neuropil_ is the surrounding neuropil signal. MLspike (https://github.com/MLspike) was used with autocalibration to estimate unitary event amplitude, decay time and noise for each region of interest and infer spiking activity. The analyses reported here used the archived discrete inferred spike counts stored in the spikes field.

### Thresholding and avalanche identification

An avalanche was defined as consecutive nonempty analysis bins bounded by empty bins after thresholding and temporal coarse-graining. For each recording and coarse-graining factor *k*, we counted complete events over a grid of population thresholds and averaged counts over all *k* temporal offsets. We fitted a Gaussian to these counts as a function of log_10_ threshold. If μ and σ denote its fitted center and width, the selected threshold was *θ* = 10^*µ* − *σ*^. The full-population analysis used 161 threshold values; subset analyses used 81.

Thresholding was applied to the original imaging frames before coarse-graining. Within each frame, neuronal contributions were sorted in ascending order, with equal values ordered independently at random for each frame. Contributions were removed while their cumulative sum remained at or below θ. The full contribution of the first neuron that crossed θ and all larger contributions were retained. Frames whose total activity did not exceed θ were set to zero. Random ties used NumPy default_rng with seed 20261027 plus the recording index, float32 random keys and 2000-frame chunks.

### Temporal coarse-graining

Temporal coarse-graining summed k adjacent thresholded frames. The original sampling interval was approximately 22 ms; *k* = 6 therefore corresponds to approximately 132 ms. Size–duration relationships were evaluated for *k* = 1–25. All *k* possible offsets were included as overlapping views of the recorded activity. Incomplete final coarse bins and events without an empty bin on both sides were excluded. Temporal offsets were not treated as independent biological observations.

### Avalanche size decomposition

For each avalanche, S is the summed inferred activity over neurons and bins, N is the number of distinct neurons with a positive contribution after population-based thresholding, and B is the number of occupied neuron–time bins. An activation is one occupied neuron–time bin after thresholding. We calculated *Ā* = *B*/*N* and *Ū* = *S*/*B*, giving the exact event-level identity

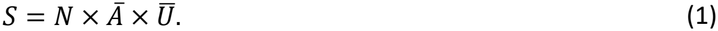

These definitions require *Ā* ≥ 1, with *Ā* = 1 for every one-bin avalanche. Exactly-once participation was the fraction of participants occupying one bin. The repeat-activation fraction was 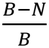, with occupied neuron–time bins as its denominator. Multiple inferred spikes within a bin contribute to *Ū*. All quantities were calculated per event before averaging. Products of separately averaged components need not equal mean *S*, and independently fitted component slopes need not sum exactly to the size exponent (Supplementary Note 2).

### Scaling fits and temporal controls

Duration-conditioned means were calculated within recordings, requiring at least 10 event views at each exact duration. We fitted unweighted ordinary least squares regressions of log mean size on log duration separately for each recording. The primary fit used L = 1–4 bins. We also fitted L = 2–4 to assess the influence of one-bin avalanches. Multibin participation summaries used events lasting at most 1 s, corresponding to *L* = 2–7 at *k* = 6. For Fig. 2c, each curve was divided by its mean size at 1 s, interpolated in log–log coordinates. This changes the vertical scale only.

The circular-shift control independently shifted each complete neuronal trace by an offset sampled uniformly over its recording length. Five shifts were analyzed per recording at *k* = 6, with the threshold refitted for each shifted population. Recorded and shifted estimates were compared within mice. The regressions describe mean size conditional on duration; they are not goodness-of-fit tests of a power-law size distribution or independent tests of a critical point.

### Correlated and random neuronal subsets

Neurons were ranked by the Pearson correlation of their inferred activity with the summed activity of all other neurons. We retained 20%, 40%, 50%, 60%, 80% or 100% of neurons, rounding the retained count upward, and evaluated *k* = 1–16. Correlation-ranked subsets used a fixed ranking within the selection segment. Random subsets were nested within each of five random permutations per recording. Thresholds were refitted for each subset and k. In the held-out comparison, neurons were ranked on one recording half and evaluated on the other; selection and evaluation halves were then exchanged. Three random permutations per half provided the held-out random comparison. The full-population condition has no subset-selection variability.

### Unbiased recruitment and comparison models

For each avalanche, 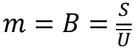 assignments were allocated independently and uniformly to *M* neurons, where *M* was the observed neuronal count for that recording. The pool size was fixed and was not fitted. The expected number of distinct participants was

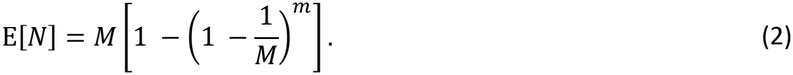

We distinguished the ratio 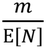 from the exact expectation 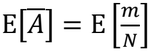. The occupancy probability *P*_*j*_ (*n*) after j assignments obeys

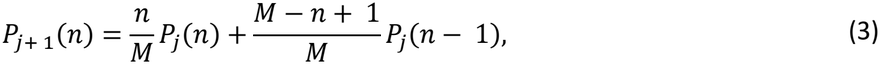

starting from 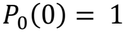. We calculated 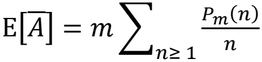. For sufficiently large assignment counts, full occupancy was used once the bound on missing any neuron fell below 10^−12^.

A within-bin comparison model sampled distinct neurons within each bin, preserving the observed active-neuron count *n*, with independent sampling across bins. Its expected recruitment was 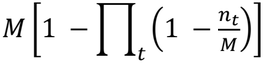. A rate-weighted model used probabilities proportional to inferred neuronal activity in the opposite recording half, giving 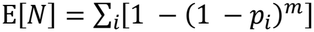. Events crossing the half-recording boundary were excluded from that comparison. All predictions and relative errors, 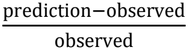, were calculated per avalanche before averaging.

### First and repeat neuronal participation

Within each avalanche, an active neuron was a first participant if it had not contributed in an earlier bin and a repeat participant otherwise. First and repeat counts sum to the number of active neurons in each bin. Activity per active neuron is the summed activity in that bin divided by its active-neuron count. Profiles were conditioned on exact duration *L* = 2–7 at *k* = 6, aligned to event onset and displayed at bin ends. Only observed active bins were used; no zero start or endpoints were added. Each recording and duration required at least 10 event views.

### Effective recruitment pool

Under uniform selection, *x* previous participants occupy a fraction 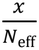 of an accessible pool. For each recording, elapsed bin and total-duration group, we estimated

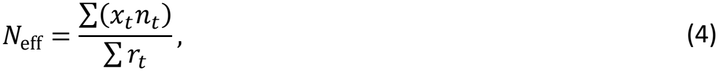

where xt counts distinct participants before bin t, nt counts currently active neurons, and rt counts current repeat participants. Sums run over event views in the same elapsed bin and total-duration group. Groups were 1–2, 2–4, 4–7 and 7–12 s, including the lower bound and excluding the upper bound. At least 10 views were required; estimates with no repeats were undefined and omitted. Estimates were not capped at the observed neuronal count. The displayed trajectory depends on the uniform-selection assumption, repeat counts and changing event composition across elapsed time.

### Statistical analysis and reproducibility

The mouse was the independent biological unit for Figures 2–4. Event-level measurements were averaged within recordings, recordings were weighted equally within each mouse, and the five mice were weighted equally.

Duration curves and profiles were conditioned on duration before this aggregation. Scalar 95% confidence intervals used the t distribution across the five mouse estimates with four degrees of freedom. Shaded bands show one standard deviation across mice. Displayed duration curves require coverage by at least three mice; exact durations are shown through *L* = 9 and logarithmic duration groups thereafter. Fits use exact durations. Overlapping temporal offsets, repeated shifts and repeated subset selections do not increase the biological sample size.

Sensitivity analyses examined threshold positions *z* = −2, −1 and 0 in *θ* = 10^*µ* + *zσ*^, alternative threshold-crossing rules, the archived continuous activity field, an independent tie seed, a fixed neuron tie order, and offset-zero-only event extraction. Two entries in the jRGECO-deepIP files, 5358193_201001 and 0013963_201023, carry a different archived indicator label; both were retained in the primary analysis and excluded in a separate sensitivity analysis. The held-out *k* = 12 subset comparison is descriptive and is not a multiple-comparison-corrected test across retained fractions or bin widths.

## Data availability

The preprocessed imaging data used for Figures 2–4 are available at https://doi.org/10.5281/zenodo.7703224. The analysis used the 17 recordings in the five jRGECO-deepIP files.

## Code availability

The published population-avalanche analysis code is available at https://github.com/PlenzLab/ParabolicAvalanches. The present analyses use an independent Python implementation of the thresholding, participation, subset, model and effective-pool procedures described in Methods.

## Acknowledgements

This research was supported by the Division of the Intramural Research Program (DIRP) of the National Institute of Mental Health (NIMH), USA, ZIAMH002797. This research utilized the supercomputing resources of the National Institutes of Health (NIH, USA; Biowulf, http://hpc.nih.gov). This manuscript was assembled and edited with assistance from Claude Code (Opus 5.5; Anthropic) and GPT-6-Astra (OpenAI). Codex was used in the reconstruction of some of the coincidence intervals in Figures 2 – 4 verified by K.S. The authors take full responsibility for the scientific content and final text.

The contributions of the authors were made as part of their official duties as NIH federal employees, are in compliance with agency policy requirements, and are considered Works of the United States Government. However, the findings and conclusions presented in this paper are those of the author(s) and do not necessarily reflect the views of the NIH or the U.S. Department of Health and Human Services.

## Author contributions

K.S., T.L.R. and D.P. conceived the study. P.K. performed data acquisition. K.S. performed the analysis and modeling. K.S., T.L.R. and D.P. wrote the manuscript.

## Supplementary Information

Ongoing activity in anatomically recurrent cortex is predominantly feedforward

### Supplementary Note 1. Expected recruitment under independent sampling

Let m be a fixed positive integer number of assignments to a fixed pool of M neurons. Assignment probabilities p_i_ are nonnegative, sum to one and remain constant across independent draws. Let N denote the number of neurons selected at least once. A neuron contributes to N unless all m draws miss it, giving

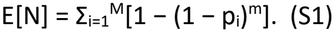

The function f(p) = 1 − (1 − p)^m^ is concave on the probability interval for integer m ≥ 1. For m > 1, its second derivative is −m(m − 1)(1 − p)^m−2^ wherever defined, with the boundary handled by continuity. Applying the concavity inequality to the mean probability gives

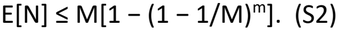

Uniform probabilities attain this upper bound. Thus, uniform sampling maximizes expected recruitment among independent assignments with fixed neuronal selection probabilities, at fixed m and M. This mathematical result does not establish equal recruitment probabilities or independence in the recorded neurons.

The maximum distinct occupancy without the independence restriction is min(m, M). A recruitment rule that avoids previously selected neurons can attain that count until the pool is exhausted. Consequently, the uniform model does not minimize repetition over all possible recruitment processes.

For a realized avalanche with m assignments and N occupied neurons, mean assignments per participant equal Ā = m/N. Averaging across independent realizations gives E[Ā] = m E[1/N]. Since the reciprocal is convex for positive N, E[Ā] ≥ m/E[N]. These quantities coincide only when N has no variability, or approximately when its fluctuations are sufficiently small. Supplementary Fig. 1b compares the exact expectation with the ratio prediction.

The relation m = S/Ū is exact when Ū is total activity divided by occupied neuron–time bins for the same avalanche. A ratio of separately averaged quantities need not reproduce an integer assignment count or the average of event-level predictions. Here, predictions use the observed integer count B for each event. Uniform independent assignments discard the temporal allocation of activations; the within-bin model separately preserves the observed active-neuron count in each bin.

### Supplementary Note 2. Exact participation constraints

For one avalanche, let B denote the number of occupied neuron–time bins after the final thresholding and coarse-graining operations, and N the number of neurons with at least one occupied bin. Every participant contributes at least one occupied bin, so B ≥ N ≥ 1. With Ā = B/N and Ū = S/B, the product N × Ā × Ū equals S, and Ā ≥ 1. Any mean of Ā with positive weights also satisfies this lower bound. A displayed mean below 1 would require a different definition or an inconsistency in counting, normalization or averaging.

Each of the N participants has exactly one first occupied bin. Thus, B − N occupied bins represent repeat participation. The fraction of occupied-bin activations that are first participations is N/B = 1/Ā; the repeat fraction is 1 − 1/Ā. These are fractions of activations, not fractions of neurons. An average repeat fraction must be computed with the stated weighting of events or occupied bins; substituting a mean Ā into the nonlinear expression generally gives a different result.

For a recorded avalanche lasting one analysis bin, B = N and Ā = 1 by definition, even if a neuron generated several action potentials within that bin. The unbiased model can assign more than one activation to the same neuron even for a one-bin avalanche. Evidence for limited repeated participation should therefore include multibin events and its dependence on temporal resolution. Coarse-graining can merge occupied bins from the same neuron, and thresholding can remove weak bins. These operations must be specified before the exact identities are compared with the plotted statistics.

### Supplementary Note 3 | Recruitment controls and model averaging

The unbiased recruitment model assigns each observed activation independently to a recorded neuron with equal probability. We compared this model with two alternatives to test whether agreement depended on ignoring the distinct-neuron constraint within a bin or differences in neuronal activity rates. The within-bin model preserves the number of distinct active neurons in each bin. The rate-weighted model estimates selection probabilities from inferred activity in the opposite recording half, excluding events that cross the split. Neither alternative reduced the mean absolute relative error in recruitment for multibin avalanches lasting up to 1 s. Thus, these comparisons do not require a more elaborate selection rule to account for the observed participation curves; they do not establish that cortical selection probabilities are equal.

For mean activations per participant, the exact model expectation averages m/N over the occupancy distribution for each event. The ratio m/E[N] is an approximation because averaging and division do not commute. Their curves nearly overlap in this dataset (Supplementary Fig. 1b). The main figure uses the exact expectation; this comparison checks that the approximation does not drive the recruitment interpretation.

**Supplementary Figure 1.**
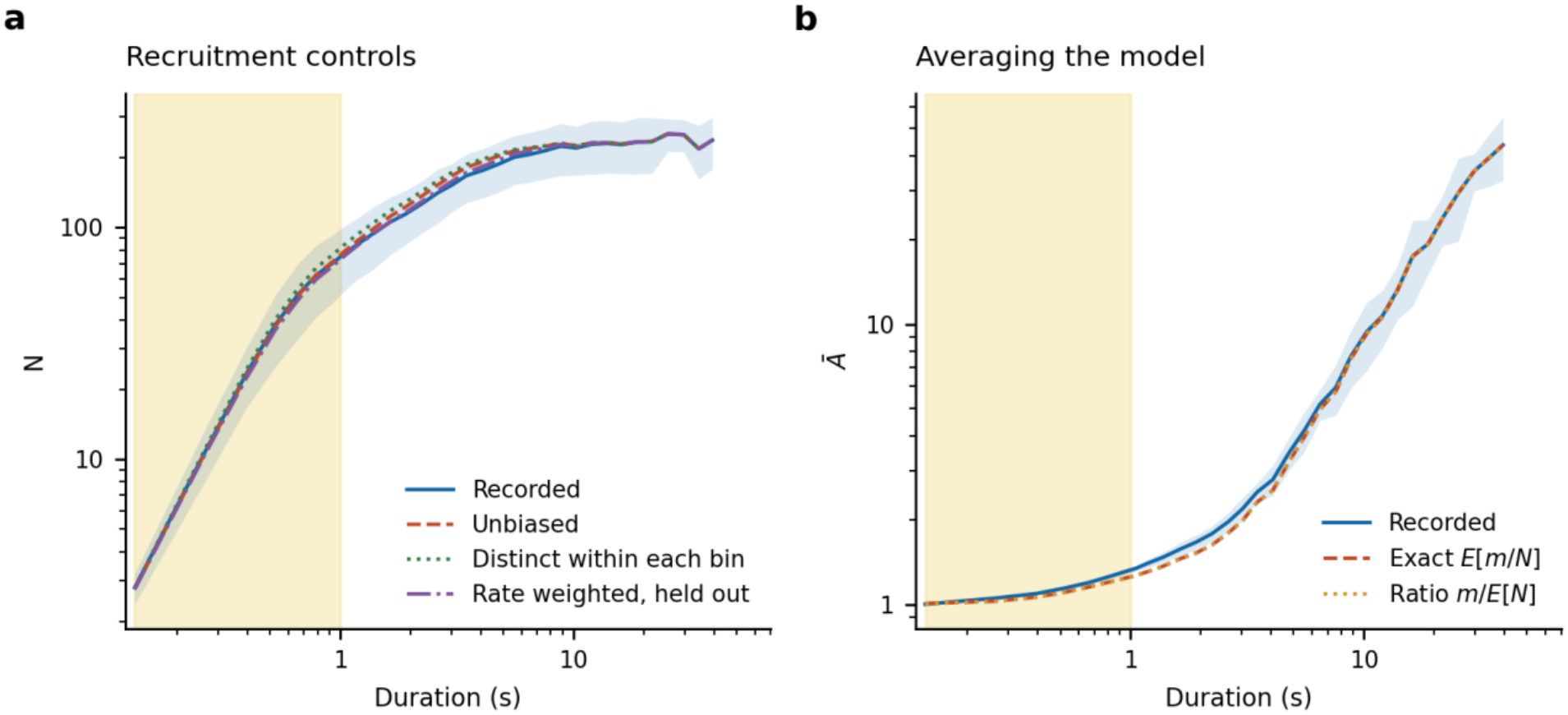
Recruitment controls and model averaging. a, Distinct participants at k = 6 in recorded activity (blue), unbiased recruitment (red dashed), the model preserving distinct active neurons within each bin (green dotted), and the rate-weighted model with probabilities estimated from the opposite recording half (purple dash-dot). b, Mean activations per participant, Ā, in the data (blue), the exact unbiased-model expectation E[m/N] (red dashed), and the ratio m/E[N] (orange dotted). All predictions use the observed event activation count m and recorded neuronal count M without fitting a pool size. Yellow shading marks durations from one analysis bin to 1 s; numerical multibin summaries exclude one-bin events. Duration bins and averaging match main Fig. 3c,d. Recording-level curves are averaged within each mouse and then across mice with equal weight. The shaded blue bands show one standard deviation across mice. Models and the exact occupancy recurrence are defined in the main Methods.

